# Neurovascular uncoupling in schizophrenia: A bimodal meta-analysis of brain perfusion and glucose metabolism

**DOI:** 10.1101/834002

**Authors:** Niron Sukumar, Priyadharshini Sabesan, Udunna Anazodo, Lena Palaniyappan

## Abstract

SUKUMAR, N., S. Priyadharshini, U. Anazodo, L. Palaniyappan. Neurovascular uncoupling in schizophrenia: A bimodal meta-analysis of brain perfusion and glucose metabolism. NEUROSCI BIOBEHAV REV X(X) XXX-XXX, XXXX. - The use of modern neuroimaging approaches has demonstrated resting-state regional cerebral blood flow (rCBF) to be tightly coupled to resting cerebral glucose metabolism (rCMRglu) in healthy brains. In schizophrenia, several lines of evidence point towards aberrant neurovascular coupling, especially in the prefrontal regions. To investigate this, we used Signed Differential Mapping to undertake a voxel-based bimodal meta-analysis examining the relationship between rCBF and rCMRglu in schizophrenia, as measured by Arterial Spin Labeling (ASL) and ^18^Flurodeoxyglucose Positron Emission Tomography (FDG-PET) respectively. We used 19 studies comprised of data from 557 patients and 584 controls. Our results suggest that several key regions implicated in the pathophysiology of schizophrenia such as the frontoinsular cortex, dorsal ACC, putamen, and temporal pole show conjoint metabolic and perfusion abnormalities in patients. In contrast, discordance between metabolism and perfusion were seen in superior frontal gyrus and cerebellum, indicating that factors contributing to neurovascular uncoupling (e.g. inflammation, mitochondrial dysfunction, oxidative stress) are likely operates at these loci. Hybrid ASL-PET studies focusing on these regions could confirm our proposition.

## 1. INTRODUCTION

Since the time of Ernst von Feuchtersleben who coined the term psychosis in 1845 (Beer, 1995), psychotic disorders have been suspected to be associated with disturbances in cerebral blood supply. This has been thoroughly investigated through the use of modern neuroimaging techniques, which have uncovered abnormalities in the resting-state regional cerebral blood flow (rCBF) across various brain regions in schizophrenia. The frontal lobe, anterior cingulate cortex, temporal lobe and occipital lobe, among others, are regions that have been showed to differ with respect to rCBF in patients compared to subjects (Oliveira et al., 2018). In healthy brains, rCBF is tightly coupled to resting cerebral glucose metabolism (rCMRglu), which increases with synaptic activity. This coupling, also known as functional hyperemia, is accomplished by the coordinated activity of a group of cells (comprised of astrocytes, endothelial cells, and neurons) called the neurovascular unit. These cells detect changes in synaptic activity, and initiate vasodilation or vasoconstriction responses to accommodate for the resultant changes in rCMRglu. (Muoio et al., 2014).

Two imaging modalities that have been very useful in studies investigating rCBF and rCMRglu in patients are Arterial Spin Labeling (ASL) and Positron Emission Tomography (PET). ASL is a relatively recent neuroimaging modality that was developed as a non-invasive analogue to gadolinium contrast MRI for the measurement of rCBF. Instead of using a potentially toxic contrast to visualize blood flow, a radiofrequency pulse is applied at the neck region to magnetize blood water molecules flowing into the brain. This allows for the capturing of a “tagged” image in the area of interest by MRI. By quantitatively comparing the tagged image with a (non-RF pulse) control image, researchers can construct an accurate representation of cerebral blood flow. (Telischak et al., 2015). Likewise, an accurate representation of rCMRglu can be constructed using PET neuroimaging. PET is a functional imaging technique that uses a radioactive tracer to measure the regional activity of the biological molecule that the tracer is attached to. A common tracer is ^18^flurodeoxyglucose (FDG), and it is often employed in neuroimaging studies to measure the cerebral metabolic rate of glucose. (Miele et al., 2008). ASL and FDG PET imaging have been used in various case-control studies to quantify case-control differences between schizophrenic patients and healthy controls.

The vascular hypothesis of schizophrenia suggests that one of the underlying mechanisms for this disorder is the disruption of the appropriate rCBF response to changes in cerebral metabolic activity (Hanson and Gottesman, 2005). Identifying regions where this uncoupling occurs is extremely important as the underlying mechanism and its pathophysiological relationship to schizophrenia can be studied in more detail. This has been done to some degree; uncoupling has been demonstrated to occur in patients with schizophrenia, especially in the prefrontal regions during task-related activities (Bachneff, 1996). However, no simultaneous ASL-PET studies identifying regions with concordance or discordance between metabolism and perfusion have been reported to our knowledge. To address this gap, we undertook a voxel-based bimodal meta-analysis to examine the relationship between rCBF and rCMRglu in schizophrenia. We hypothesized that several brain regions would show combined abnormalities of perfusion and metabolism, while uncoupling of these two parameters will be observed in prefrontal regions. The meta-analysis was performed using the Anisotropic Effect Size version of Seed-based d Mapping (AES-SDM) and only included studies that have used the Montreal Neurological Institute (MNI) or Talairach coordinate system to report findings. AES-SDM was also used to conduct meta-regression analyses to quantify the effect of nuisance variables such as age, gender, duration of illness, antipsychotic dosage, and illness severity on the heterogeneity of the findings.

## 2. METHODS

### 2.1. ASL Study Quality

The methodological quality of the included studies was assessed by scoring 4 criteria encompassing all aspects of data collection and analysis which can confound the quality of the ASL data and subsequently bias the validity and reliability of the study inferences. The 4 criteria are: 1) participant selection, 2) image acquisition, 4) image pre-processing and analysis, and 5) statistical analysis techniques. The criteria for appropriate quality standards for image acquisition were based on the recommended implementations of ASL by the ISMRM Perfusion Group and the European ASL in Dementia (Alsop et al., 2015). The minimum standard requirement for image acquisition were modified since 36% of the included ASL studies were completed 2-3 years after the recommended guidelines. Emphasis was placed on determining whether the included studies specified imaging parameters that impact the signal-to-noise in ASL images. ASL image quality increases as the signal-to-noise increases. The criteria were scored as either ‘adequate’, if all aspects of the criteria were reported and met minimum standard, or ‘inadequate’, if aspects of the criteria were missing or does not meet minimum standard, or ‘unclear’, if no clear conclusion could be drawn from the information provided. These criteria are further detailed in Figure 3.

**Figure 1:**
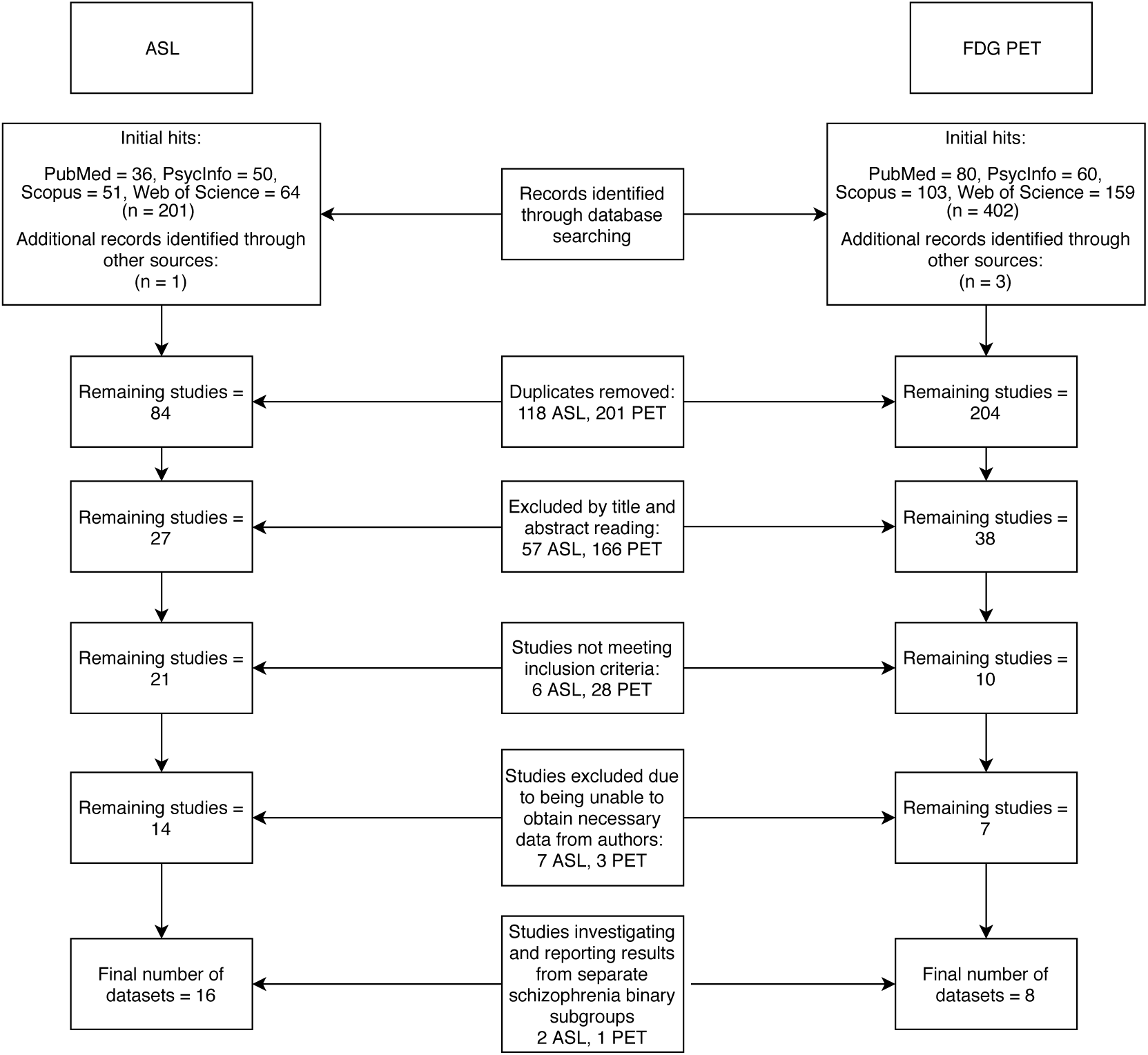
Flowchart of literature search.

**Figure 2:**
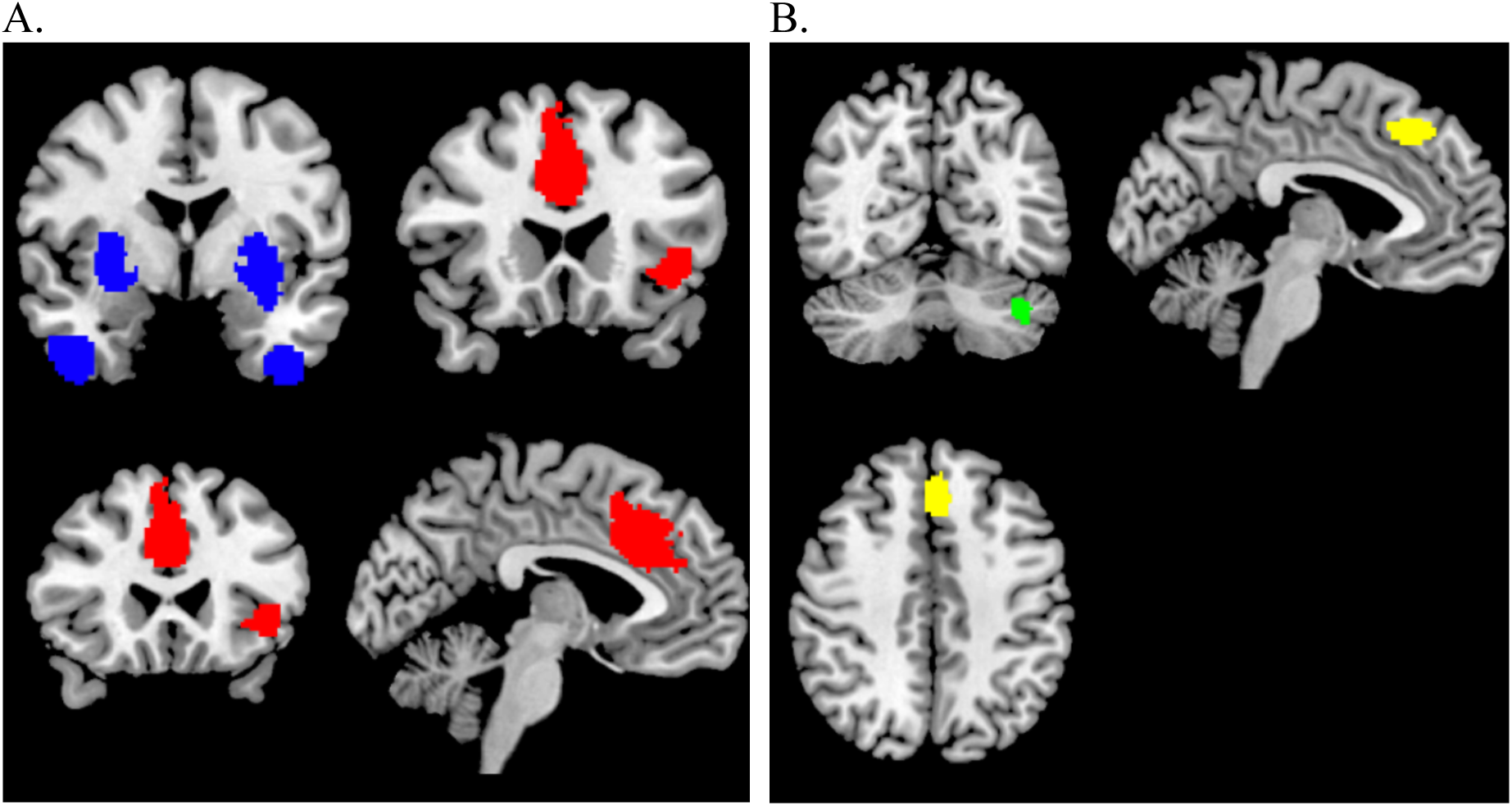
Regions of conjoint findings are shown on the left (a), and regions with disjoint findings are shown on the right (b). Bilateral striatum and temporal pole were found to have conjoint increases in rCBF and rCMRglu (shown in blue). Left frontoinsular cortex and bilateral dorsal anterior cingulate cortex were found to have conjoint reductions in rCBF and rCMRglu. Regional neurovascular uncoupling was notable in the left superior frontal gyrus (−6,30,44; Left BA 8; SDM-z = -2.001, p=0.00033, reduced rCMRglu, normal rCBF – shown in yellow) and left cerebellum (−38,-66,-34; crus I and II; SDM-Z=2.27, p=0.00026, no. of voxels = 129; increased rCMRglu, normal rCBF – shown in green).

**Figure 3:**
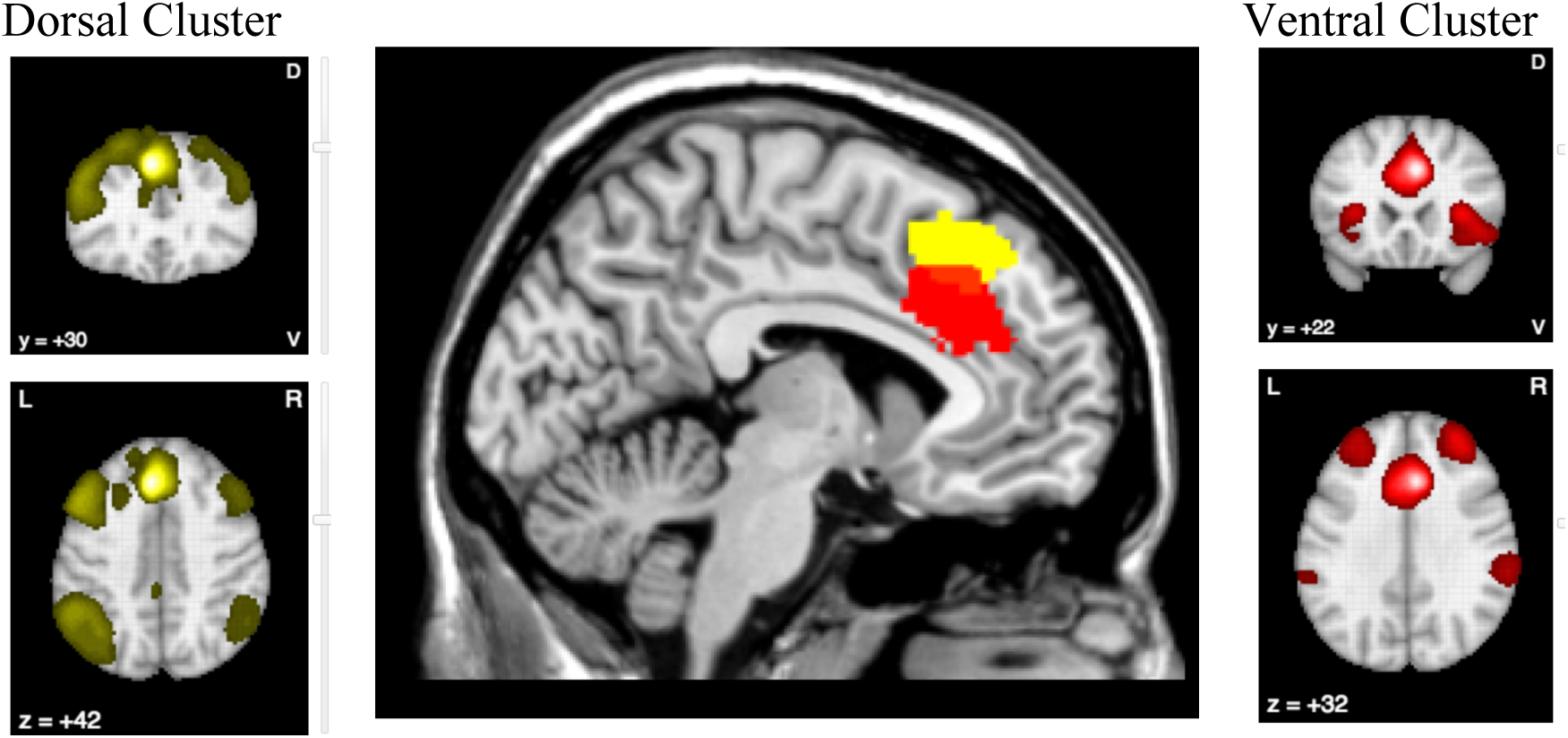
Anterior cingulate region with conjunction and disjunction findings belong to different brain networks. Using Neurosynth functional connectivity meta-analytical database, the two peak coordinates are plotted to show the network level differences. The more ventral cluster (red) has reduced rCMRglu as well as reduced rCBF while the more dorsal (yellow; -6,30,44) has normal rCBF despite reduced metabolism. The ventral cluster is well connected to the Salience Network, while the dorsal node seems to participate in the frontoparietal executive network.

### 2.2. PET Study Quality

The methodological quality of the included studies was assessed by scoring 5 criteria encompassing all aspects of data collection and analysis which can confound the quality of the PET data and subsequently bias the validity and reliability of the study inferences. The 5 criteria are: 1) participant selection, 2) participant preparation, 3) image acquisition, 4) image pre-processing and analysis, and 5) statistical analysis techniques. The criteria for appropriate quality standards for participant preparation and image acquisition were based on the SNMMI Procedure guideline for FDG PET Brain Imaging Version 1.0 (Waxman et al., n.d.). The criteria were scored as either ‘adequate’, if all aspects of the criteria were reported and met minimum standard, or ‘inadequate’, if aspects of the criteria were missing or does not meet minimum standard, or ‘unclear’, if no clear conclusion could be drawn from the information provided. These criteria are further detailed in Figure 4.

**Figure 4:**
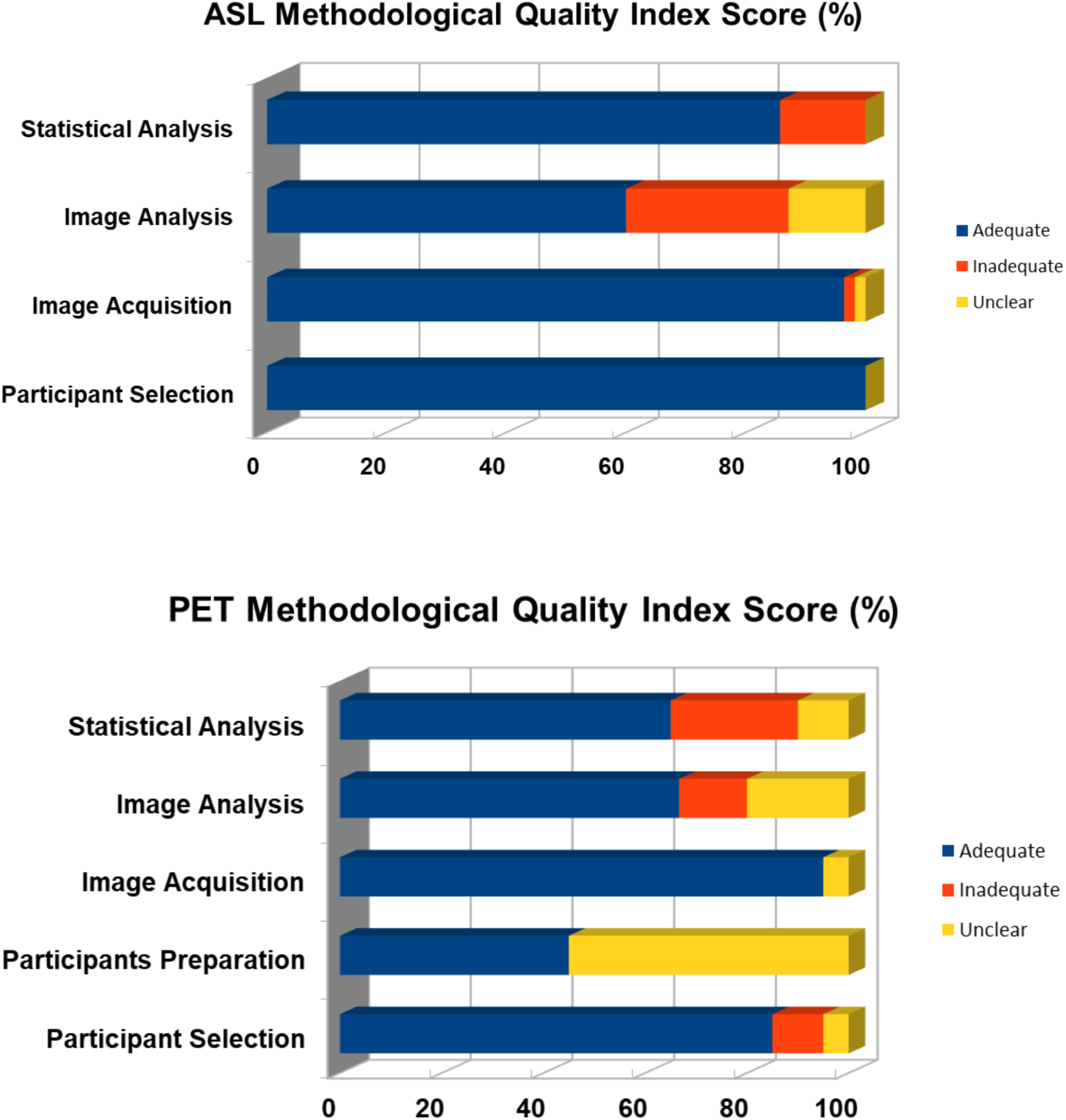
The Quality Index Score is the proportion of studies that are scored as adequate, inadequate, or unclear.

### 2.3. Search

Two literature searches were conducted across four databases (PubMed, PsycInfo, Scopus and Web of Science). The search terms ‘arterial’, ‘spin’, ‘labeling’, and ‘schizophrenia’ yielded 83 results, and the terms ‘FDG’, ‘PET’ and ‘schizophrenia’ yielded 201 results (after duplicates were removed). The following inclusion criteria was applied: Case control studies reporting voxelwise ASL or FDG-PET changes in schizophrenia using MNI or Talairach. Studies that met the inclusion criteria, but did not report all of the data required for the meta-analysis were not immediately excluded in the hope that a correspondence with the authors of these studies could be initiated to obtain the missing data. Studies that did not use ICD 10 or DSM IV/5 diagnostic criteria for schizophrenia or did not investigate the whole brain were excluded. 21 ASL and 8 PET papers remained after inclusion and exclusion criteria were applied to the search results. Of these, 12 papers did not report their data in a format that was compatible with our meta-analysis. We contacted the authors of these papers and were only able to obtain additional data from two studies (Oliveira et al., 2018), (Liu et al., 2012). Three of the papers we finally included in our meta-analyses (Walther et al., 2017), (Pinkham et al., 2015), (Ben-Shachar et al., 2007), investigated schizophrenia binary subgroups (for example, catatonic and non-catatonic schizophrenia patients) compared to controls. The final meta-analyses thus used 16 datasets from 14 ASL papers and 6 datasets from 5 PET papers to generate results.

### 2.4. AES SDM Analysis

AES-SDM creates meta-analytic maps of studies that use MNI or Talairach coordinates to denote brain regions of significant group differences, weighted by sample size, variance, and between-study heterogeneity The software was used to generate a meta-analytic map for the compiled datasets, using permutation tests (p<0.001, cluster extent k=10) to determine statistical significance of results. The sensitivity of reported findings to exclusion of individual studies was assessed by performing a jackknife analysis. This was done by repeating the meta-analysis multiple times, each time leaving out one of the studies that was originally included it. A score was given to each reported brain region corresponding to the number of times it was reported in the meta-analyses. The effect of the following variables was studied by conducting a meta-regression analysis: age, gender, duration of illness, anti-psychotic dosage, and illness severity. The meta-regression analyses were thresholded conservatively (P < 0.0005) to minimize false correlations as a result of the limited variability of some of these factors.

### 2.5. Conjunction and Disjunction Analysis of rCBF and rCMRglu Changes

We used multimodal analysis to identify which of the brain regions identified in our meta-analysis showed significant changes in both CBF and rCMRglu. Our goal was not necessarily to demonstrate a physiological correlation between these measures, but rather to identify which of our results were supported by data from both ASL and PET imaging studies. As this was a 4-tailed test (allowing 2 directions of results in 2 modalities), we used a conservative threshold of p<0.0025 for cluster inclusion and p<0.00025 (10-times more stringent) for peak identification, based on the minimum acceptable threshold for conjunction in each individual modality map as p<0.05. This method, described in detail by Radua et al. (Radua et al., 2013), has been used in various studies for bimodal conjunction meta-analysis (Wise et al., 2016), (Radua et al., 2012), (Radua et al., 2014). We also used the meta-regression procedure and tested the slope (1m0) of the effect of the measure (ASL vs PET) on effect-sizes reported in the SDM, to identify brain regions showing changes consistent with neurovascular uncoupling (altered rCBF with normal rCMRglu or altered rCMRglu with normal rCBF). We used MRIcron software to generate visual representations of conjunction and disjunction findings.

### 2.6. Meta-regression analysis

We explored the influence of age, gender, duration of illness, PANSS total symptom severity and overall dose of antipsychotic medications (in chlorpromazine equivalents) on the reported effect-sizes. To reduce spurious relationships, in line with prior studies (Radua et al., 2012) we used a probability threshold of 0.0005, and tested the slope (1m0) (e.g. comparing effect sizes in studies with lowest vs. highest values of the predictor variable of interest), and restricted meta-regression to findings detected in the main analyses (Radua and Mataix-Cols, 2009). We also visually inspected the regression plots from peak coordinates and discarded slopes driven by <5 studies (Radua et al., 2012).

## 3. RESULTS

### 3.1. Study Demographics

Our meta-analysis drew data from 22 datasets from 19 studies. In total, 557 patients with schizophrenia were compared to 584 healthy controls. Table 1 lists the demographic data for the participants in each study. Most studies published number of patients and healthy controls, patient gender distribution, patient age, duration of illness, antipsychotic dosage, syndrome severity, type of scan, and study quality. Almost all the studies matched healthy controls to patients for at least age and gender, with some studies matching controls for other variables such as level of education. Duration of illness, antipsychotic dosage and PANSS scores ranged from 25.2 (SD 5.58) to 92.0 (SD 22) for ASL studies and from 33.8 (SD 5.3) to 84.7 (SD 23.6) for PET studies. With a few exceptions, all included ASL studies used a 3T scanner with a pCASL technique to obtain results and only included patients diagnosed with schizophrenia specifically. One study (Cui et al., 2017) used a pASL technique, one study (Horn et al., 2009) used a 1.5T MRI scanner and three studies (Kindler et al., 2018), (Kindler et al., 2015), (Liu et al., 2012), included individuals with schizoaffective disorder in their patient groups. All PET studies only included patients diagnosed with schizophrenia disorder specifically. The authors from studies with missing clinical information were contacted, and some of the unpublished data was retrieved for this review.

**Table 1:** Demographic table for datasets included in meta-analysis. DOI, duration of illness (years). PANSS, Positive and Negative Syndrome Scale. CLPZ Eq., chlorpromazine equivalents.

| Study | Modality | Patients | Controls | Patient Age | Control Age | % Females | PANSS | Duration of Illness | CLPZ eq | Quality scores |
| --- | --- | --- | --- | --- | --- | --- | --- | --- | --- | --- |
| Kindler 2018 | ASL | 32 | 31 | 41.6 ± 13.4 | 39.4 ± 12.3 | 43.75 | 76.6 ± 17.4 | 14.3 | 497 ± 210 | 0.8 |
| Walther 2017 (Catatonia) | ASL | 15 | 41 | 35.9 ± 12.7 | 38.6 ± 13.6 | 26.67 | 77.1 ± 18.9 | 12.8 ± 12.0 | 461.3 ± 346.4 | 0.53 |
| Walther 2017 (Non-Catatonia) | ASL | 27 | 41 | 37.1 ± 10.6 | 38.6 ± 13.6 | 37.04 | 67.1 ± 16.3 | 10.5 ± 10.6 | 373.6 ± 359.4 | 0.53 |
| Pinkham 2015 (Paranoid) | ASL | 16 | 25 | 38.5 ± 7.71 | 33.64 ± 12.42 | 50 | 32 ± 5.37 | NA | 332.2 ± 546.75 | 0.93 |
| Pinkham 2015 (Non-Paranoid) | ASL | 16 | 25 | 38.8 ± 13.24 | 33.64 ± 12.42 | 31.25 | 25.2 ± 5.58 | NA | 350.5 ± 571.26 | 0.93 |
| Zhu 2015 | ASL | 100 | 94 | 33.6 ± 8.6 | 33.3 ± 10.4 | 43 | 71.3 ± 22.7 | 10.2 ± 8.2 | 453.2 ± 342.9 | 0.73 |
| Kindler 2015 | ASL | 34 | 27 | 41.5 ± 12.9 | 39.8 ± 12.6 | 47.06 | 75.5 ± 17.3 | NA | 518.9 ± 235.7 | 0.73 |
| Ota 2014 | ASL | 36 | 42 | 37.9 ± 12.6 | 37.9 ± 13.0 | 52.78 | 61.8 ± 19.3 | 16.8 ± 11.3 | 604.8 ± 459.2 | 0.73 |
| Pinkham 2011 | ASL | 30 | 24 | 35.7 | 35.73 | 40 | NA | NA | 373.7 | 0.93 |
| Walther 2011 | ASL | 11 | 14 | 35.3 ± 12.54 | 31.71 ± 6.08 | 27.27 | 54.2 ± 14.11 | 8.9 ± 13.29 | 442.5 ± 241.03 | 1.00 |
| Scheef 2010 | ASL | 11 | 25 | 32 | 30 | 27.27 | 43.1 ± 8.5 | NA | NA | 0.93 |
| Horn 2009 | ASL | 13 | 13 | 29.6 ± 11.2 | 26.6 ± 4.6 | 38.46 | 63.9 | 2.79 ± 2.6 | 556.2 | 0.87 |
| Oliveira 2018 | ASL | 28 | 28 | 32.8 ± 7.8 | 31.1 ± 5.8 | 14.29 | 69.4 ± 17.2 | 14.3 | 546.4 | 1.00 |
| Cui 2017 (Non-AVH) | ASL | 25 | 25 | 24 ± 5 | 26 ± 5 | 48 | 92 ± 22 | 1.5 ± 1.75 | NA | 0.87 |
| Stegmayer 2017 | ASL | 20 | 30 | 38.2 ± 11.4 | 36.7 ± 12.9 | 61.7 | 72.6 ± 17.1 | 12.2 ± 12.3 | 400.2 ± 344.2 | 1.00 |
| Liu 2012 | ASL | 19 | 20 | NA | NA | 57.90 | NA | 20.5 ± 10.0 | 622.1 ± 418.0 | 0.67 |
| Park 2009 | PET | 29 | 21 | 29.8 ± 3.9 | 3.9 ± 3.0 | 48.27 | 33.8 ± 4.8 | 7.5 ± 4.1 | 633.2 ± 415.6 | 0.65 |
| Desco 2002 | PET | 51 | 18 | 31.93 | 13.16 | 31.37 | NA | 7.89 | NA | 0.6 |
| Kim 2017 | PET | 19 | 18 | 32.6 ± 12.0 | 30.7 ± 7.9 | 36.84 | 84.7 ± 23.6 | 2.3 ± 1.8 | NA | 0.75 |
| Ben-Shachar 2006 (HPS) | PET | 8 | 8 | 35.9 ± 11.8 | 34.5 ± 11.5 | 37.5 | 43.7 ± 6.4 | 15.57 ± 11.3 | NA | 0.65 |
| Ben-Shachar 2006 (LPS) | PET | 8 | 8 | 42.7 ± 15.3 | 34.5 ± 11.5 | 50 | 33.6 ± 5.3 | 18.42 ± 10.3 | NA | 0.65 |
| Horga 2014 | PET | 9 | 8 | NA | NA | NA | 45 ± 8.2 | 2.1 ± 7.8 | NA | 0.8 |

### 3.2. Study Quality

The average quality scores for the ASL and PET studies used in the meta-analysis were 0.84 and 0.69 respectfully. Among the ASL studies, the lowest quality score was 0.67 (Liu et al., 2012) and among the PET studies the lowest quality score was 0.6 (Ben-Shachar et al., 2007). The Quality Index Scores for these studies are visually represented in Figure 4.

### 3.3. Pooled differences across the 2 modalities

The SDM analysis showed several brain regions of significant difference in rCBF or rCMRglu between patients and healthy controls. These results are listed in Table 2. Patients had significantly increased rCBF or rCMRglu in the right lenticular nucleus (putamen, BA 48), left striatum, right inferior temporal gyrus (BA 20), left temporal pole (middle temporal gyrus, BA 20), right thalamus, and corpus callosum, and significantly reduced rCBF or rCMRglu in the right median cingulate (paracingulate gyri, BA 24), right middle occipital gyrus (BA 18), left inferior frontal gyrus (triangular part, BA 47), left superior occipital gyrus (BA 17), and right superior frontal gyrus (dorsolateral, BA 8). Other than the right superior frontal gyrus, which only survived 17 cross-validations, each of the other regions survived at least 21 cross-validations. The right putamen, left striatum, right inferior temporal gyrus, right median cingulate, and right middle occipital gyri survived all 22 cross-validations. Finally, meta-regression analysis was conducted to investigate the relationship between the change in neurological activity in these regions and various nuisance variables of interest. Antipsychotic dosage was found to be correlated to changes in various brain regions. These relationships are graphically represented in Figure 3. No other variables were found to have any significant correlations.

**Table 2:** Brain regions of significant difference in rCBF or rCMRglu between patients with schizophrenia and controls. Jack-knife analysis was scored out of 22. Voxel threshold: P < 0.005; Peak height threshold: peak SDM-Z > 1.000; Extent threshold: cluster size ≥ 10 voxels.

| Region | SDM Z | MNI Coordinate | Cluster Size | P value | Jack-knife Score | Cluster breakdown |
| --- | --- | --- | --- | --- | --- | --- |
| **Patients>controls**<br>Blobs of $\geq 59$ voxels with all voxels SDM-Z $\geq 1.655$ and all peaks SDM-Z $\geq 1.982$ | | | | | | |
| Right Putamen | 3.517 | 28,4,12 | 1411 | $< 0.0001$ | 22 | BA48<br>BA34 |
| Left Striatum | 3.186 | -22,2,6 | 1080 | $< 0.0001$ | 22 | BA48<br>BA34 |
| Right Inferior Temporal Gyrus | 3.186 | 42,2,-42 | 596 | $< 0.0001$ | 22 | BA20<br>BA36 |
| Left Temporal Pole | 2.470 | -34,0,-44 | 295 | 0.0005 | 21 | BA20<br>BA36 |
| Right Anterior Thalamic Projections | 2.204 | 18,-22,12 | 103 | 0.0005 | 21 |  |
| Corpus Callosum | 1.982 | 18,-28,4 | 59 | 0.001 | 21 | n/a |
| **Controls>patients**<br>Blobs of $\geq 101$ voxels with all voxels SDM-Z $\leq -1.894$ and all peaks SDM-Z $\leq -2.218$ | | | | | | |
| Right Median Cingulate Gyrus | -2.829 | 8,26,32 | 1292 | 0.0005 | 22 | BA24<br>BA32 |
| Right Middle Occipital Gyrus | -2.873 | 30,-92,12 | 357 | $< 0.0001$ | 21 | BA18<br>BA19 |
| Left Superior Occipital Gyrus | -2.641 | -16,-100,14 | 274 | $< 0.0001$ | 22 | BA17<br>BA18 |
| Left Inferior Frontal Gyrus, (Triangular) | -2.708 | -42,22,-2 | 647 | $< 0.0001$ | 21 | BA47<br>BA45<br>BA48 |
| Right Superior Frontal Gyrus, (Dorsolateral) | -2.218 | 20,20,56 | 101 | 0.001 | 17 | BA8 |

**Table 3:** Quality Analysis Tool for included ASL studies.

| Each criterion was categorized as follows:<br>Adequate: All aspects of the criteria were reported and met minimum standard.<br>Inadequate: Aspects of the criteria were missing or did not meet minimum standard.<br>Unclear: No clear conclusion could be drawn from the information provided. |  |  |
| --- | --- | --- |
|  | Criteria | Minimum standard |
| 1 | Participant Selection | I. Participants in the study should be subjects with a diagnosis of schizophrenia or schizoaffective disorder established through clinical chart review using structured diagnostic interviews based on DSM-IV/V and healthy controls between ages of 18 and 65 years.<br>II. Participants were selected following a prospective inclusion and exclusion criteria.<br>III. Details should be provided about age distribution, female to male ratio and disease description as minimum information (subject demographics table).<br>IV. Total number of subjects per group were >10. |
| 2 | Image acquisition | *Emphasis placed on factors that impact signal-to noise.*<br>I. Hardware considerations: Scanner manufacturer and Field Strength specified was 1.5 T or 3T.<br>II. The specific ASL pulse labelling approach was specified. The four labelling approaches are: continuous (CASL), pseudo-continuous (pCASL), pulsed (PASL), and velocity-selective (VS-ASL).<br>III. Time between labeling and imaging (post-label delay) was specified<br>IV. Total imaging time was specified, or enough information was specified to estimate total imaging time (repetition time, echo time, and the number of label and control pairs). |
| 3 | Image analysis | I. Measures to restrict motion or correct motion was specified. *ASL is sensitive to motion and motion can reduce signal-to-noise in ASL.*<br>II. The following parameters used in calculating CBF must be specified:<br>The blood-brain partition coefficient used<br>The T1 of blood used<br>The Labelling efficiency used<br>III. The use of a brain atlas or template for spatial normalization was specified. |
| 4 | Statistical analysis | I. Statistical methods for controlling multiple comparisons were specified for significant differences.<br>II. The influence of age and gender was controlled/removed. *\*\* Age and gender are significant CBF modifiers (Clement et al JCBFM 2017)* |

**Table 4:** Quality Assessment Tool for included PET studies.

|  | **Criteria** | **Minimum standard** |
| --- | --- | --- |
| 1 | Participant Selection | V. Participants in the study should be subjects with a diagnosis of schizophrenia or schizoaffective disorder established through clinical chart review using structured diagnostic interviews based on DSM-IV/V and healthy controls between ages of 18 and 65 years.<br>VI. Participants were selected following a prospective inclusion and exclusion criteria.<br>VII. Details should be provided about age distribution, female to male ratio and disease description as minimum information (subject demographics table).<br>VIII. Total number of subjects per group were >10. |
| 2 | Participant preparation | V. Participants fasted for 4-6 hours.<br>VI. The use of caffeine or alcohol, substance use/abuse/dependence, or psychoactive medications that may affect cerebral glucose metabolism were specified.<br>VII. The blood glucose was checked prior to FDG-injection and was no greater than 150 – 200 mg/dl.<br>VIII. Participants were kept in a stable environment during PET uptake period. This includes a quiet, dimly-lit room and eyes open/closed. |
| 3 | Image acquisition | IV. Scanner make and model was specified.<br>V. Administered FDG dose was within 185-740 Mbq<br>VI. Uptake period within 20-60 minutes<br>VII. Emission scan duration of a minimum of 15 minutes |
| 4 | Image analysis | III. An attenuation correction method was specified.<br>IV. The reconstruction algorithm was specified.<br>V. Measures to restrict motion or correct motion was specified.<br>VI. The use of a brain atlas or template for spatial normalization was specified.<br>VII. An appropriate reference region selection (cerebellar cortex or whole brain) for count normalization was reported. |
| 5 | Statistical analysis | I. Exclusion of none gray matter voxels was reported.<br>II. Statistical methods were specified for significant differences<br>III. The influence of age and gender was controlled/removed. |

### 3.4. Conjunction and Disjunction Analyses

Among patients with schizophrenia, we observed a conjoint reduction in rCBF and rCMRglu in the right median cingulate / paracingulate gyri and left inferior frontal gyrus. A conjoint increase in rCBF and rCMRglu was noted in the right putamen and right inferior temporal gyrus. (Voxel probability threshold: p=0.0025, Peak height threshold: p=0.00025, Cluster extent threshold: 10 voxels). Regional neurovascular uncoupling was notable in the superior frontal gyrus (reduced rCMRglu, normal rCBF) and cerebellum (increased rCMRglu, normal rCBF). These regions are visually represented in Figure 2.

## 4. DISCUSSION

To our knowledge, this is the first bimodal neuroimaging meta-analysis which combines information from whole brain FDG-PET studies investigating resting metabolic state, and ASL studies investigating the resting blood flow in schizophrenia. We report 3 major findings: (1) frontoinsular cortex and bilateral dorsal anterior cingulate cortex show reduced rCBF as well as rCMRglu in schizophrenia (2) bilateral dorsal striatum and temporal pole show increased rCBF as well as rCMRglu in schizophrenia (3) brain regions with rCBF changes consistently show rCMRglu changes, but brain regions with rCMRglu changes are not always coupled with rCBF changes, especially in superior frontal (dorsomedial ACC) and cerebellar cortices.

The frontoinsular cortex and bilateral dorsal anterior cingulate cortex showing reduced rCBF as well as rCMRglu in schizophrenia belong to the Salience Network. Many prior studies have extensively investigated the connectivity within the SN and between SN and other brain regions, and have highlighted the concentration of grey matter reduction in schizophrenia around the nodes of the SN. Our findings once again highlight the primary role that the SN plays in the diagnostic construct of schizophrenia. The neural basis of insula-related observations in fMRI studies have been considered with some caution given the vascular anatomy of this region. Our bimodal analysis establishes that defects in glucose utilization of SN nodes are a key aspect of the pathophysiology of schizophrenia. We also note that the frontoinsular cortex shows a notable reduction in rCBF/rCMRglu in patients with longer duration of illness, indicating the possibility of a progressive defect in this key node.

Studies enrolling patients on higher doses of antipsychotics are associated with larger rCBF/rCMRglu effects in the bilateral dorsal striatum. The effect of antipsychotics on striatal rCBF in schizophrenia has been studied extensively, and our results are highly consistent with the synthesis reported by Goozée et al. (Goozée et al., 2014). Striatal D2 blockade seems to have a direct effect on increasing metabolism as well as blood flow of striatum, while other brain regions do not exhibit a relationship of similar magnitude. Increased rCBF/rCMRglu of temporal pole is consistent with medial temporal lobe pathology that has been reported in the literature (Gur et al., 2000), (Lee et al., 2016), (Crespo-Facorro et al., 2004). Given that the posited function of the temporal pole is one of emotion-perception binding (Crespo-Facorro et al., 2004), hypermetabolism in this region may be related to acute psychotic symptoms as shown by Crespo-Feccorro et al., though our meta-analysis is not able to confirm or refute this notion.

We observed rCBF/rCMRglu uncoupling in 2 sites: dorsomedial prefrontal cortex and cerebellar vermis. The presence of normal rCBF in a site with reduced rCMRglu indicates a relative hyperperfusion, and resonates with the observation made by Taylor et al (Taylor et al., 1999) who demonstrated that ACC region demonstrates a relative hyperperfusion in subjects with schizophrenia. Coupling of CMR and CBF appears to be linked to a vasoactive mechanism, as well as a structural factor related to capillary density (Kuschinsky et al., 1981). Postmortem studies in schizophrenia have not uncovered any notable vascular structural pathology to date, except for an increase in astrocytic end-feet in the prefrontal cortex shown by one study (Uranova et al., 2010). In certain mitochondrial encephalopathies and lactic acidosis, uncoupling of rCBF and CMRglu occurs (Shishido et al., 1996), (Sano et al., 1995), indicating that alterations in oxidative stress pathways may be relevant to our current observation in schizophrenia.

There are several limitations in this review that should be considered when interpreting results. Firstly, the cerebellar disjunction findings must be considered with caution as the quality of ASL signals from this region has been suboptimal in many studies. The lack of cerebellar coverage as well as the influence of ASL labeling site on posterior cerebral circulation may have influenced the reported disjunction. Secondly, all of the reported PET studies used CT-based anatomical registration, wile ASL uses MR-based information. This might have influenced the exact location of peak coordinates, though the spatial smoothing used in SDM mitigates this to some extent. We also urge caution in interpreting the negative results from meta-regression (i.e. lack of age, gender and severity effects on metabolism) as none of the individual studies were powered to detect these relationships, and the meta-regression approach cannot deal with non-linear effects. Similarly, disjunction results could also be driven by the well-known issue of false negative results from coordinates based meta-analyses (see Albajes-Eizagirre & Radua, 2018 for a review). Finally, our results do not demonstrate that metabolic and vascular abnormalities are necessarily correlated at the subject (or group) level. Our aim was more modest, and restricted to localizing those brain regions where both abnormalities co-exist in schizophrenia.

To conclude, schizophrenia related regional perfusion abnormalities capture the aberrant metabolism of underlying neuro-glial tissue. In specific brain regions, such as the dorsomedial prefrontal cortex, neurovascular uncoupling suggestive of possible inflammation (causing inappropriate hyperemia), astroglial dysfunction or mitochondrial defect are likely to be present. This uncoupling needs further characterization, possibly using hybrid PET/MRI, to establish a mechanistic basis. These observations raise an interesting question of whether focused pharmacological restoration of blood-flow regulation could alleviate symptoms of schizophrenia.

## DISCLOSURES

LP reports personal fees from Otsuka Canada, SPMM Course Limited, UK, Canadian Psychiatric Association; book royalties from Oxford University Press; investigator-initiated educational grants from Janssen Canada, Sunovion and Otsuka Canada outside the submitted work. PS reports personal fees from Otsuka Canada, SPMM Course Limited, UK, outside the submitted work. All other authors report no relevant conflicts.

## ACKNOWLEDGEMENTS

This study was supported by The Chrysalis Foundation. LP acknowledges support from the Tanna Schulich Chair of Neuroscience and Mental Health and the Opportunities Fund of the Academic Health Sciences Centre Alternative Funding Plan of the Academic Medical Organization of Southwestern Ontario (AMOSO).

## APPENDIX

## SUPPLEMENTARY MATERIAL

**Table S1.** rCBF or rCMRglu abnormalities in schizophrenia: meta-regression analyses

|  | Talairach<br>Coordinates | SDM<br>z-value<br>(a) | P Value<br>(b) | No. of<br>voxels<br>(c) | Breakdown<br>(No. of voxels)<br>(c) |
| --- | --- | --- | --- | --- | --- |
| <u>EFFECTS OF ILLNESS DURATION</u> |  |  |  |  |  |
| *Decreased rCBF or rCMRglu in patients with longer illness duration (high < low duration)* |  |  |  |  |  |
| Left inferior frontal gyrus (triangular) | -52,42,-2 | -2.118 | 0.00001 | 125 | Left BA45<br>Left BA46<br>Left BA47 |
| <u>EFFECTS OF ANTIPSYCHOTIC MEDICATION</u> |  |  |  |  |  |
| *Increased rCBF or rCMRglu in patients receiving higher doses of antipsychotics (high > low doses)* |  |  |  |  |  |
| Left striatum | -22, 0, 2 | 1.912 | <0.0001 | 586 | Left BA48 |
| Right striatum | 32,-12,-4 | 1.489 | 0.00011 | 139 | Right BA48 |

(a) Voxel probability threshold: $p = 0.0005$ (difference in slope)
(b) Peak height threshold: $z = 1$
(c) Cluster extent threshold: 10 voxels. Regions with less than 10 voxels are not reported in the cluster breakdown.

**Table S2.**
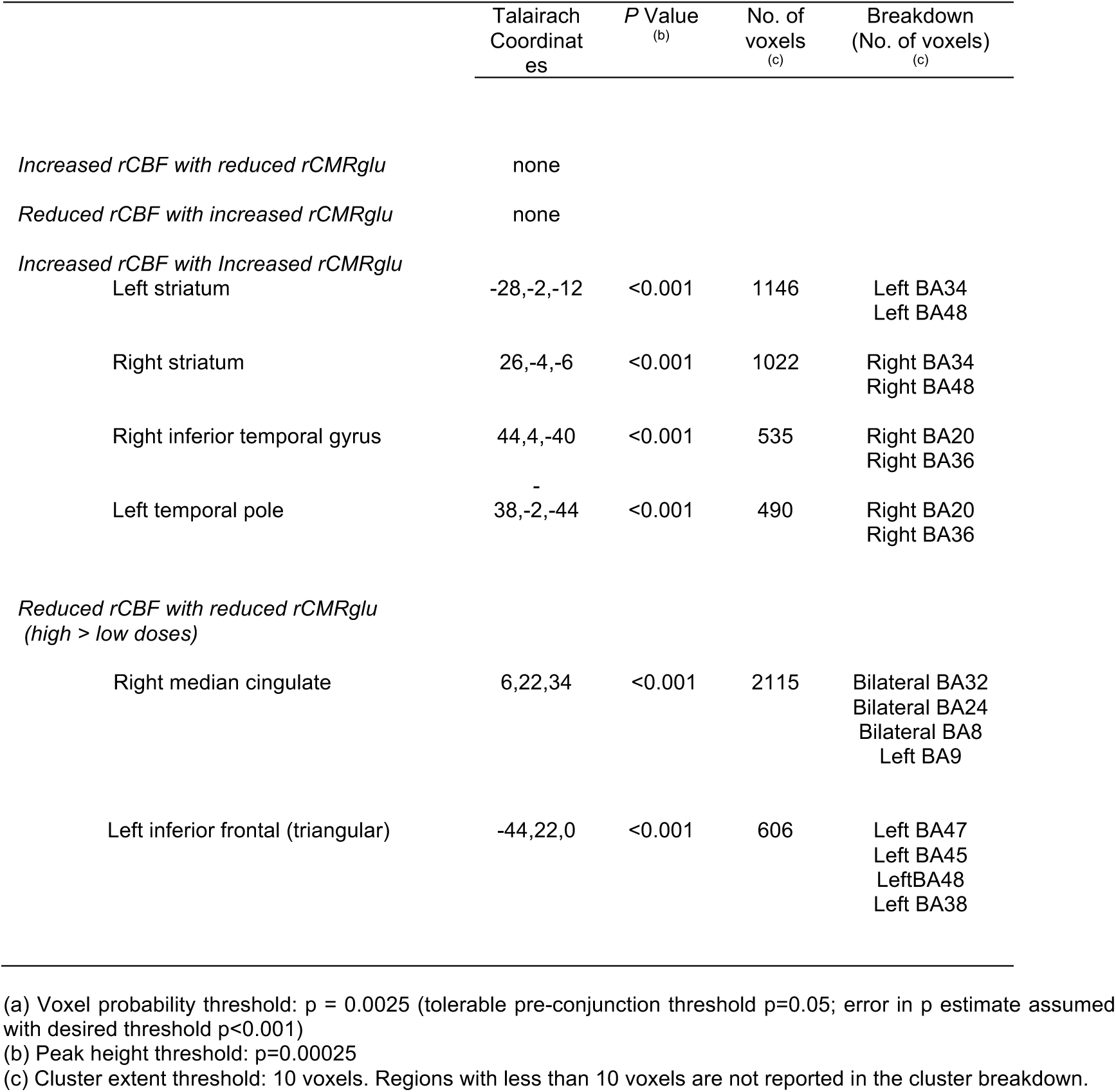
Combined rCBF and rCMRglu abnormalities in schizophrenia: multimodal analyses

